# Estimation of heritabilities and genetic correlations by time slices using predictivity in large genomic models

**DOI:** 10.1101/2023.06.28.546953

**Authors:** Ignacy Misztal

**Affiliations:** Animal and Dairy Science, University of Georgia, Athens, GA 30602, USA

**Author notes:** **Corresponding author:** Ignacy Misztal, Animal and Dairy Science, University of Georgia, Athens, GA 30602.

**Keywords:** genetic parameters, estimation, genomic selection, correlated response

## Abstract

Under genomic selection, genetic parameters may change rapidly from generation to generation. Unless genetic parameters used for a selection index are current, the expected genetic gain may be unrealistic, possibly with a decline for antagonistic traits. Existing methods for parameter estimation are computationally unfeasible with large genomic data. We present formulas for estimating heritabilities and genetic correlations applicable for large models with any number of genotyped individuals. Heritabilities are calculated by combining 2 formulas for genomic accuracies: one that relies on predictivity and another that depends on the number of independent chromosome segments. Genetic correlations are calculated from predictivities across traits. Both formulas include approximate standard errors. The formula for heritabilities was evaluated based on information for 4 commercial data sets extracted from published studies. Calculated heritabilities were close to those used initially, except for a much lower new heritability for one single case; that heritability was later confirmed as valid. Formulas for genetic correlations were tested with simulated data and 1,000 replicates. The formula-based estimates were always close to the values assumed for simulation. Standard errors were high with 1,000 validation animals but small with 10,000. The proposed formulas can be used routinely as a check on the evaluation system whenever the number of validation individuals is large enough. If excessive changes are detected from generation to generation, the selection index can be modified appropriately.

## INTRODUCTION

Selection causes a change in genetic parameters dependent on the intensity of selection (Falconer and Mackay 1996; Walsh and Lynch 2018). Before genomic selection, various studies on animal populations indicated declining heritabilities and more antagonistic correlations (e.g. Holm *et al*. 2004; Haile-Mariam and Pryce 2015).. The heritability of a growth trait in pigs halved after a few years of genomic selection, whereas its correlation with a fitness trait became 50% more antagonistic (Hidalgo *et al*. 2020). That study used slices of data because of computing limitations, and the computation took months.

Accurate genetic parameters are important for realistic trait trends and setting up a selection index (Hazel 1943; Hazel *et al*. 1994). With incorrect parameters, trends may be exaggerated and lead to unrealistic estimates of genetic gain. Also, increased antagonism between production and fitness traits with old parameters may lead to the decline of important fitness traits. If such declines are not addressed early by a change in the selection index, a reversal may take a long time because of the low heritabilities of fitness traits.

Variance components are routinely estimated with methods that have good theoretical properties, such as restricted maximum likelihood (REML) or Bayesian via Gibbs sampling (Sorensen and Gianola 2002). Such methods rely on the sparseness of mixed model equations and can accommodate large data sets without genomic information (Misztal 2008). With genomic information, mixed models are no longer sparse, which leads to expensive computation. With specialized software that accommodates dense blocks in sparse matrices, REML was applicable with up to 40k genotyped individuals for single-trait models and less with multiple-trait models (Masuda *et al*. 2015; Misztal *et al*. 2021). Many commercial data sets across species now have over 200k genotyped individuals, with over 6 million for U.S. dairy cattle (https://uscdcb.com/milestones).

Current methods estimate component variances for a base population. With possibly rapid changes in genetic parameters, estimating parameter changes in time slices would be useful. This study investigates methods that can estimate heritabilities and genetic correlations with any number of genotyped individuals and for any time slice.

## METHODS

### Predictivity and theoretical accuracies

Legarra *et al*. (2008) showed that the average accuracy of an evaluation (*acc*) for selected animals could be calculated from predictivity, a correlation between their adjusted phenotypes and estimated breeding values:

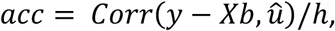

where *y* − *Xb* is a vector of observations adjusted for fixed effects for the selection candidates, 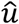 is EBVs for these animals calculated with their observations removed from the dataset, and h^2^ is heritability. Daetwyler *et al*. (2008) proposed a formula for *acc* in a population with *M*_*e*_ independent chromosome segments, *N* genotyped and phenotyped animals, and heritability *h*^2^:

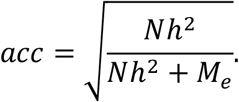

If both formulas are applicable and estimates of *M*_*e*_ are available (e.g. from Pocrnic *et al*. 2017), the estimate of heritability can be derived by equating both formulas:

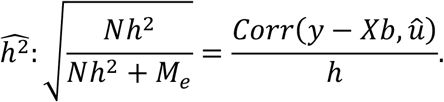

Then,

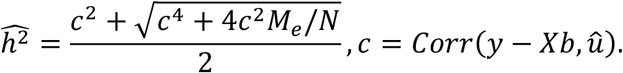

### Predictivity across traits and genetic correlations

The predictivity formula can be generalized to estimate genetic correlations. Assume correlated traits *i* and *j* with additive variances 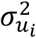 and 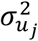, respectively, and covariance 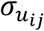. Based on the conditional distribution

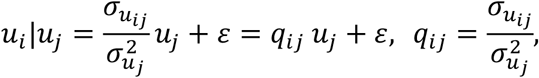

the predictivity of the adjusted phenotype of trait *i* for EBV of trait *j* is

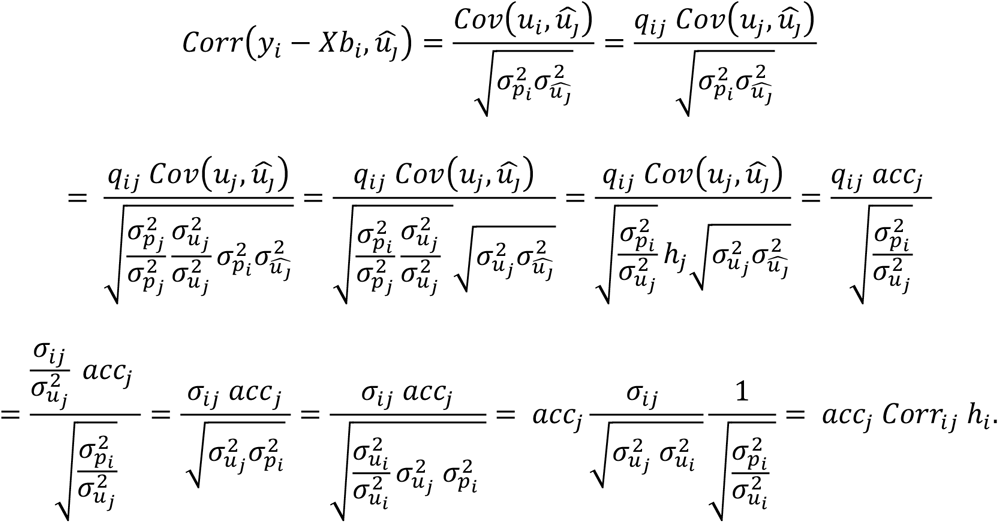

After rearranging, the genetic correlation between traits *i* and *j* is:

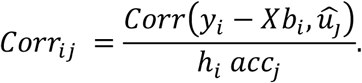

Because *Corr*_*ij*_ = *Corr*_*ji*_, asymptotically

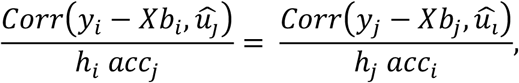

although the standard errors may differ.

### Standard errors

Under normality, the standard error of correlation *ρ* is 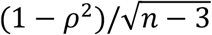, where *n* is the number of observations (Gnambs 2022). For large *n* and small *ρ*, the formula can be simplified to 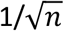. The standard error in the accuracy formula then is

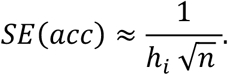

Assume a trait with a heritability of 0.25 and 1,600 observations. Then the standard error for the accuracy estimate would be 0.05 with a 95% confidence interval of about 0.2. The error would decrease with a larger validation population and a higher heritability.

The standard error in the genetic correlation formula can be calculated by treating *h*_*i*_ and *acc*_*j*_ as constants. Then

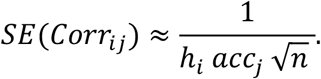

Assume 2 traits with heritabilities of 0.25, EBVs with accuracies of 0.5, and 1,600 animals in the validation population. Then the standard error would be 0.10 with a 95% confidence interval of about 0.4. With a larger validation population, higher heritability of the first trait, and higher accuracy of the second trait, the error would decrease. Note that the standard error generally would be different for *Corr*_*ij*_ and *Corr*_*ji*_, and it would be smaller for a combination with the higher denominator.

An approximate standard error for approximation of heritability can be calculated based on linear approximation:

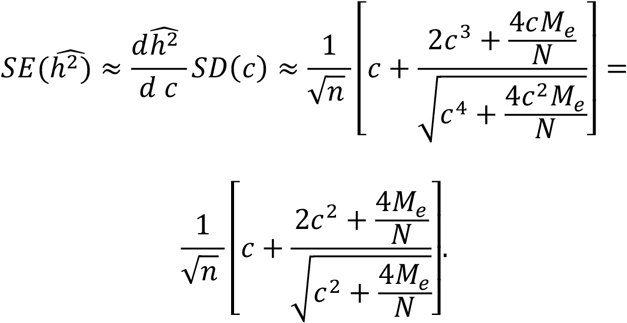

For very large datasets with *N* >> *M*_*e*_, the standard error simplifies to

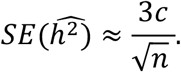

Note that *N* is the number of individuals in the reference population and *n* is the number of individuals in the validation population.

### Data

The formula for heritability was tested with data extracted from several studies, which included milk yield in Holsteins (Cesarani *et al*. 2021), growth and fitness traits in pigs (Hollifield *et al*. 2021), and a growth trait in broiler chicken (Hidalgo *et al*. 2021). The information extracted included initial heritabilities used in the calculations, approximate number of genotyped animals with phenotypes in both reference and validation populations, and calculated accuracy or predictivity. In all cases, up to 3 generations of animals were included for pig and chicken data and animals born after the year 2000 for Holsteins. The number of animals in the accuracy formula was assumed to be the number of genotyped animals with phenotypes except for fitness traits of pigs, where fewer animals had phenotypes than genotypes; for those traits, the number of animals with phenotypes was used.

Several data sets were simulated to test the formula to estimate genetic correlations using an algorithm that allowed for many replicates. Two sets of heritabilities were included: one for medium heritability traits (0.2/0.3) and one for medium and low heritability traits (0.2/0.02). Various sizes of validation sets were included. The algorithm steps were:

1. Generate a vector of breeding values *i*, (*i* = 1, … *n*) from a bivariate distribution:

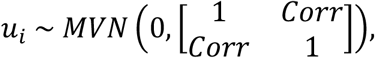

where the correlation between traits (*Corr*) ranged from −0.5 to 0.5 and *n* was 1k or 10k.
2. Generate uncorrelated residuals from a bivariate distribution such that when combined with the breeding values would result in expected heritabilities:

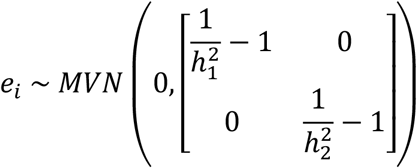

for heritabilities of 0.3, 0.2, and 0.02.
3. Generate observations:

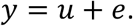
4. Calculate estimated breeding values with the desired accuracy (*<u>acc</u>*):

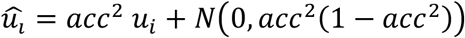

for *acc* of 0.8 and 0.3.

### Analyses

For heritability calculation, extracted data were directly used in the formula for heritability when predictivity was available. If only computed accuracies were available from the studies, those were converted to predictivity using the formula predictivity = *acc* × *h*_*comp*_, where *h*_*comp*_ was the computed heritability from the study. The number of independent chromosome segments followed the empirical results of Pocrnic *et al*. (2017) and were 15k for Holsteins and 5k for pigs and chickens.

For simulated data to test the formula to estimate genetic correlations, computations included direct computation of accuracy as 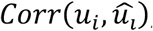, accuracy by predictivity, predictive ability across traits, and estimates of genetic correlations. Standard errors were computed from 1,000 replicates and then from the analytical formula.

## RESULTS

### Heritability calculations

Table 1 shows details of heritability calculations for Holsteins and 2 traits in pigs and chickens. Holsteins had the highest number of animals with phenotypes and genotypes (580k) and the highest number of validation animals (381k). The heritability calculated from the formula was 0.33, which is close to the initial heritability of 0.35 from the study of Cesarani *et al*. (2021). The next largest dataset was for a growth trait of chickens and included about 100k animals with genotypes and phenotypes. Although Hidalgo *et al*. (2021) assumed a heritability of 0.30 based on earlier studies, the calculated heritability was lower (0.14). Discussions with the data providers indicated that based on their recent studies, a heritability of 0.13 is currently used (Vivian Breen, 2022, personal communication). For pigs, a higher heritability growth trait and a lower heritability fitness trait were examined. The initial heritability was 0.21 for the growth trait and 0.05 for the fitness trait (Hollifield *et al*. 2021), and the calculated heritability was slightly higher (0.26) for the growth trait and almost the same (0.06) for the fitness trait.

**Table 1.** Description of real data sets for estimating heritabilities

|  | Species/trait |  |  |  |
| --- | --- | --- | --- | --- |
|  | Holsteins/<br>milk | Pigs/<br>fitness | Pigs/<br>growth | Chicken/<br>growth |
| Number of animals with<br>phenotypes and genotypes<br>in reference population ($N$) | 580k | 12k<br>(generations 5–7) | 20k<br>(generations 5–7) | ~100k |
| Number of animals in<br>validation population ($n$) | 381k | 2k<br>(generation 8) | 11k<br>(generation 8) | ~40k |
| Initial heritability | 0.35 | 0.05 | 0.21 | 0.30 |
| Assumed $M_e$ | 15k | 5k | 5k | 5k |
| Calculated accuracy |  | 0.40 | 0.80 | 0.58 |
| Predictivity | 0.55 | 0.09 | 0.36 | 0.31 |
| Calculated heritability | 0.33 | 0.06 | 0.26 | 0.14 |

### Calculation of genetic correlations

Table 2 shows intermediate results and final estimates of genetic correlations for 4 simulated datasets. For all datasets, accuracy of EBVs calculated directly and by predictivity matched the simulated accuracy. All genetic correlation estimates matched, too. However, standard error depended on the volume of data, heritabilities, and correlation order. In particular, for the first 3 datasets, the standard error for prediction based on the correlation of the first trait’s phenotype with the second trait’s EBV was twice as large as for the reverse combination. In the analytical formula for standard error of the genetic correlation, the standard error is inversely proportional to the square root of heritability for the trait with a phenotype multiplied by the accuracy for the trait with the EBV. Those products are 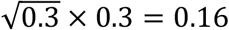 for the first combination and 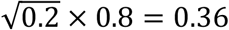 for the reverse combination. Thus, for traits with roughly similar heritability, standard error is lower for a trait with EBV of higher accuracy.

**Table 2.**
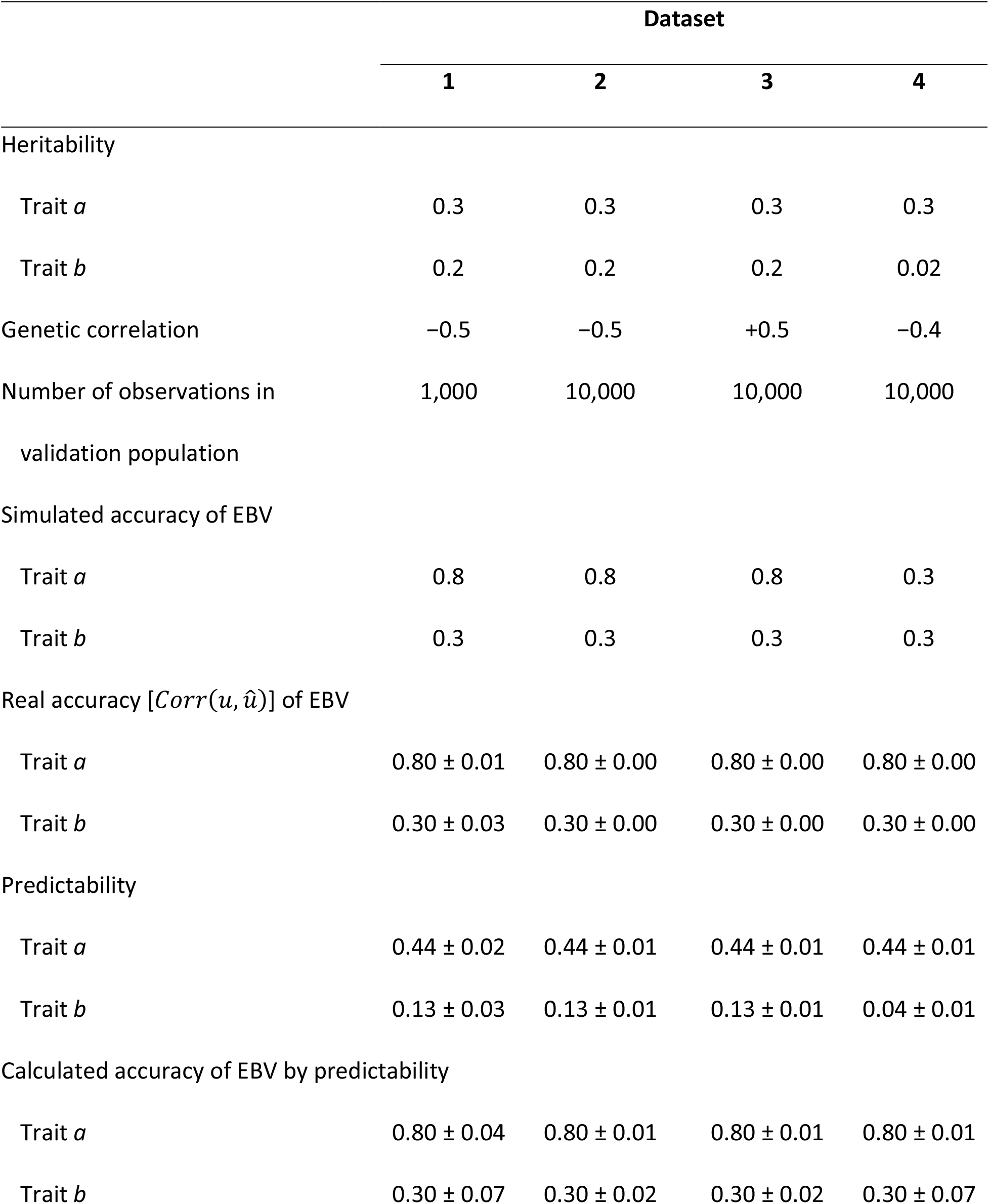
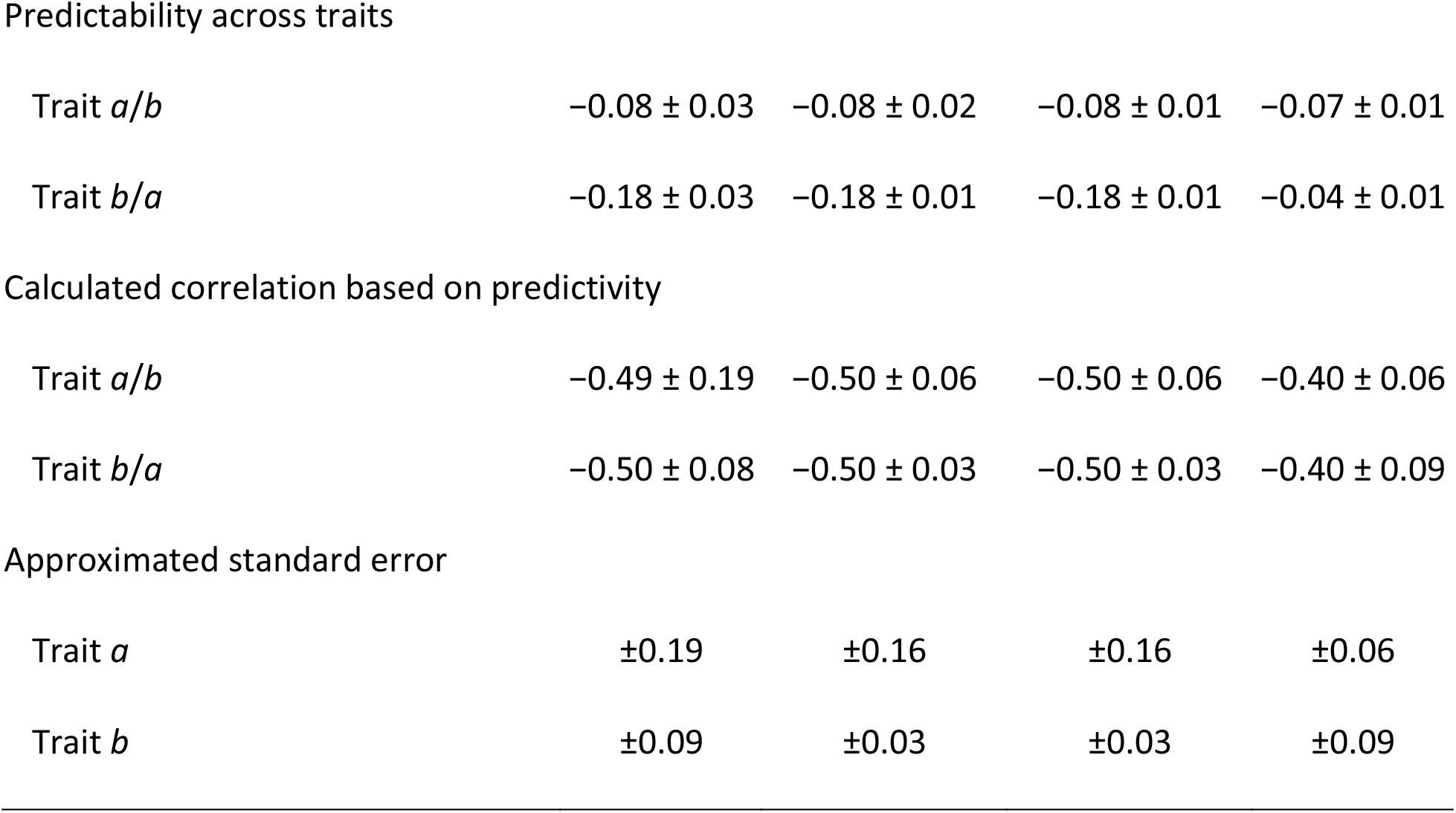
Description of simulated data sets and measures of accuracy and genetic correlations

|  | Dataset |  |  |  |
| --- | --- | --- | --- | --- |
|  | 1 | 2 | 3 | 4 |
| Heritability |  |  |  |  |
| Trait $a$ | 0.3 | 0.3 | 0.3 | 0.3 |
| Trait $b$ | 0.2 | 0.2 | 0.2 | 0.02 |
| Genetic correlation | -0.5 | -0.5 | +0.5 | -0.4 |
| Number of observations in validation population | 1,000 | 10,000 | 10,000 | 10,000 |
| Simulated accuracy of EBV |  |  |  |  |
| Trait $a$ | 0.8 | 0.8 | 0.8 | 0.3 |
| Trait $b$ | 0.3 | 0.3 | 0.3 | 0.3 |
| Real accuracy [$Corr(u, \hat{u})$] of EBV | | | | |
| Trait $a$ | $0.80 \pm 0.01$ | $0.80 \pm 0.00$ | $0.80 \pm 0.00$ | $0.80 \pm 0.00$ |
| Trait $b$ | $0.30 \pm 0.03$ | $0.30 \pm 0.00$ | $0.30 \pm 0.00$ | $0.30 \pm 0.00$ |
| Predictability |  |  |  |  |
| Trait $a$ | $0.44 \pm 0.02$ | $0.44 \pm 0.01$ | $0.44 \pm 0.01$ | $0.44 \pm 0.01$ |
| Trait $b$ | $0.13 \pm 0.03$ | $0.13 \pm 0.01$ | $0.13 \pm 0.01$ | $0.04 \pm 0.01$ |
| Calculated accuracy of EBV by predictability |  |  |  |  |
| Trait $a$ | $0.80 \pm 0.04$ | $0.80 \pm 0.01$ | $0.80 \pm 0.01$ | $0.80 \pm 0.01$ |
| Trait $b$ | $0.30 \pm 0.07$ | $0.30 \pm 0.02$ | $0.30 \pm 0.02$ | $0.30 \pm 0.07$ |
| Predictability across traits |  |  |  |  |
| Trait $a/b$ | $-0.08 \pm 0.03$ | $-0.08 \pm 0.02$ | $-0.08 \pm 0.01$ | $-0.07 \pm 0.01$ |
| Trait $b/a$ | $-0.18 \pm 0.03$ | $-0.18 \pm 0.01$ | $-0.18 \pm 0.01$ | $-0.04 \pm 0.01$ |
| Calculated correlation based on predictivity |  |  |  |  |
| Trait $a/b$ | $-0.49 \pm 0.19$ | $-0.50 \pm 0.06$ | $-0.50 \pm 0.06$ | $-0.40 \pm 0.06$ |
| Trait $b/a$ | $-0.50 \pm 0.08$ | $-0.50 \pm 0.03$ | $-0.50 \pm 0.03$ | $-0.40 \pm 0.09$ |
| Approximated standard error |  |  |  |  |
| Trait $a$ | $\pm 0.19$ | $\pm 0.16$ | $\pm 0.16$ | $\pm 0.06$ |
| Trait $b$ | $\pm 0.09$ | $\pm 0.03$ | $\pm 0.03$ | $\pm 0.09$ |

The standard errors for medium heritability traits are high with 1k validation animals (0.08 for the reverse combination of traits, which had the smaller standard errors) but reasonably small with 10k animals (0.03 for the reverse combination). Standard error increased with smaller heritability even with 10k validation animals (0.06 for the first combination). Therefore, the proposed method should be applied only to large models with a large validation population. Standard errors from the simulation by replication and standard errors calculated by the formulas were nearly identical. If standard errors from 2 combinations are markedly different, an obvious solution is to choose the estimate with the smaller analytical standard error. If both standard errors are similar, an average of both correlations can be used.

## DISCUSSION

The proposed methods create the possibility for an inexpensive estimation of heritability and genetic correlation trends by time intervals. For each interval, calculate predictivity within traits and across traits. Assuming the number of independent chromosome segments, calculate heritabilities and then genetic correlations and their standard errors. For genetic correlations, choose the estimate with the smaller standard error. The trends of genetic parameters would allow extrapolating changes and modifying the selection index to avoid future problems.

The advantage of the presented formulas is low computing costs for any model with any number of genotyped individuals as the only computationally intensive step is the genetic evaluation. However, many questions can be raised about the robustness of the formulas. The formula of Daetwyler *et al*. (2007) was derived assuming only animals with genotypes and phenotypes and under ideal conditions: no genotyping errors and, indirectly, an unlimited number of markers. Therefore, in practical cases, it may give an inflated estimate of accuracy. Field data usually have additional information from ungenotyped animals, which is considered in ssGBLUP (Aguilar *et al*. 2010), for example. Still, the genotyping strategy may be suboptimal, and both genotyping and pedigree errors exist. Subsequently, the accuracy would be smaller than in idealized conditions. When the number of genotyped animals is much larger than *M*_*e*_, a compromise choice for *N* would be the number of animals with both genotypes and phenotypes.

Another value required by the formula of Daetwyler *et al*. (2007) is *M*_*e*_. Such a number can be derived in several ways: for example, 4*N*_*e*_*L*, where *N*_*e*_ is effective population size and *L* is length of genome (Stam 1980), or the number of eigenvalues that explain 98% of variation in the genomic relationship matrix (Pocrnic *et al*. 2016). Both studies suggest an *M*_*e*_ of 5k to 15k for farm animals, and unpublished results suggest less than 2k for crosses of inbred lines in plants. With a large number of genotyped individuals, the change in accuracy in the formula of Daetwyler *et al*. (2007) with increased *N* or *M*_*e*_ is small. Assuming that having many individuals corresponds to ≥4 *M*_*e*_, the formula of heritability can be applied with reference populations of at least 20k to 60k genotyped individuals for animals, about 8k for plants, and 1.5 million for humans (*N*_*e*_ ≥ 3000).

A serious concern is the susceptibility of the predictivity to selection bias (Bijma 2012). A substantial bias was observed in earlier studies with limited genotyping (Lourenco *et al*. 2015). In a recent study (Abdollahi-Arpanahi *et al*. 2022), accuracies by predictivity and Method LR (Legarra and Reverter 2018) considered resistant to selection bias were nearly identical, which indicates that selection biases with the predictivity formula are low with widespread genotyping. The formulas for the genetic correlations could possibly be incorporated into the LR method to take advantage of simpler computing and better statistics. For example, the adjustment for added effects needs to be done explicitly in the predictivity formula, but such an adjustment is automatic in Method LR.

’In general, using established methods with good theoretical properties like REML or Bayesian via Gibbs sampling (Sorensen and Gianola 2002) is preferable. Although those methods are prohibitively expensive with large models, computing refinements may make them applicable. However, they have weaknesses aside from computing costs in some cases. First, they are susceptible to extreme selection bias with selective genotyping. In selective genotyping, where young animals are mass-selected for growth and only the best are genotyped, the estimate of heritability without genomic information is about 0.3 but reaches 0.8 with genomic information (Wang *et al*. 2020). Predictivity is not subjected to a strong bias in this case, as it only involves comparisons between phenotypes and EBVs obtained without those phenotypes. Another issue with REML and Bayesian via Gibbs sampling is sensitivity to genotyping errors. For example, using lower-density SNP information leads to lower prediction error variance and higher estimated accuracies in best linear unbiased prediction, because larger differences between genomic and pedigree information due to fewer SNPs are interpreted as added information. With lower-density SNP information, predictivity would decline. Finally, the established methods would have to be modified to estimate parameters by time slices.

## CONCLUSIONS

Genetic heritabilities and correlations can be estimated in large genomic models for any slice of data by using predictability within and across traits. Computing requirements are minimal. The standard error of the estimates depends on the size of the validation population and is reasonably small for a population of 10,000 individuals.

## ACKNOWLEDGEMENTS

This study was partially funded by Agriculture and Food Research Initiative Competitive Grant no. 2020-67015-31030 from the U.S. Department of Agriculture’s National Institute of Food and Agriculture. Useful comments and editing by Mary Kate Hollifield are gratefully acknowledged.

